# Pharmacologic Therapy for Engraftment Arrhythmia Induced by Transplantation of Human Cardiomyocytes

**DOI:** 10.1101/2021.02.15.431108

**Authors:** Kenta Nakamura, Lauren E. Neidig, Xiulan Yang, Gerhard J. Weber, Danny El-Nachef, Hiroshi Tsuchida, Sarah Dupras, Faith A. Kalucki, Anu Jayabalu, Akiko Futakuchi-Tsuchida, Daisy S. Nakamura, Silvia Marchianò, Alessandro Bertero, Melissa R. Robinson, Kevin Cain, Dale Whittington, Hans Reinecke, Lil Pabon, Björn C. Knollmann, Steven Kattman, R. Scott Thies, W. Robb MacLellan, Charles E. Murry

## Abstract

**Background:** Engraftment arrhythmias (EAs) are observed in large animal studies of intramyocardial transplantation of human pluripotent stem cell-derived cardiomyocytes (hPSC-CMs) for myocardial infarction. Although transient, the risk posed by EA presents a barrier to clinical translation.

**Objectives:** We hypothesized that clinically approved antiarrhythmic drugs can prevent EA-related mortality as well as suppress tachycardia and arrhythmia burden.

**Methods:** hPSC-CM were transplanted into the infarcted porcine heart by surgical or percutaneous delivery to induce EA. Following a screen of antiarrhythmic agents, a prospective study was conducted to determine the effectiveness of amiodarone plus ivabradine in preventing cardiac death and suppressing EA.

**Results:** EA was observed in all subjects, and amiodarone-ivabradine treatment was well-tolerated. None of the treated subjects experienced the primary endpoint of cardiac death, unstable EA or heart failure compared to 5/8 (62.5%) in the control cohort (hazard ratio 0.00; 95% confidence interval, 0–0.297; p = 0.002). Overall survival including two deaths in the treated cohort from immunosuppression-related infection showed borderline improvement with treatment (hazard ratio 0.21; 95% confidence interval, 0.03–1.01; p = 0.05). Without treatment, peak heart rate averaged 305 ± 29 beats per min (bpm), whereas in treated subjects peak daily heart rate was significantly restricted to 185±9 bpm (p = 0.006). Similarly, treatment reduced peak daily EA burden from 96.8 ± 2.9% to 76.5 ± 7.9% (p = 0.003). Antiarrhythmic treatment was safely discontinued after approximately one-month of treatment without recrudescence of arrhythmia.

**Conclusions:** The risk of engraftment arrhythmia following hPSC-CM transplantation can be reduced significantly by combined amiodarone and ivabradine drug therapy.

**Condensed Abstract:** Engraftment arrhythmia (EA) is a transient but serious complication of cardiac remuscularization therapy. Using a porcine model of cardiac remuscularization and EA, ivabradine and amiodarone were independently effective in suppressing tachycardia and arrhythmia, respectively. Baseline amiodarone combined with adjunctive ivabradine successfully prevented cardiac death, unstable EA and heart failure (hazard ratio 0.00; 95% confidence interval, 0–0.297; p = 0.002) and significantly suppressed peak daily heart rate and arrhythmia burden (p=0.006 and 0.003, respectively). Antiarrhythmic treatment was successfully discontinued after one-month without recrudescence of arrhythmia. We conclude that EA can be suppressed by combined amiodarone and ivabradine drug therapy.

## Introduction

Ischemic heart disease including myocardial infarction (MI) and heart failure remains the leading cause of death in the United States and around the world (1). Approximately one billion cardiomyocytes are permanently lost during MI (2) and an increasing proportion of MI survivors—an estimated 20 to 40% (3,4)—later develop heart failure. Current treatments can slow the initiation and progression of heart failure, but none replaces lost myocardium, short of orthotopic heart transplantation, which remains restricted in availability and indication (5). Human pluripotent stem cells (hPSCs, comprising embryonic stem cells [ESCs] and their reprogrammed cousins, induced pluripotent stem cells [iPSCs]) are a renewable source of cardiomyocytes (CMs). Transplantation of hPSC-derived cardiomyocytes (hPSC-CMs) into infarcted myocardium of small animals—mice, rats, and guinea pigs—has shown stable engraftment (6–10). More recently, our group and others have shown remuscularization and functional benefit in infarcted non-human primates (NHP) following transplantation of pluripotent stem cell-derived cardiomyocytes (11–13). In addition to functional remuscularization, the human graft vascularizes and electromechanically couples with the host myocardium within one-month post-transplant and remains durable up to three months, the longest time tested.

Although no arrhythmias were observed in smaller animals, we and others consistently observe ventricular arrhythmias following hPSC-CM transplantation in NHPs (11–13) and pigs (14) which we have called “engraftment arrhythmias” (EAs). EAs are generally transient, occurring within a week of transplantation and typically resolve spontaneously after approximately one-month. Based on electrical mapping, overdrive pacing, and cardioversion studies, EAs appear to originate focally in the graft or peri-graft myocardium and function as automatous foci rather than reentrant pathways (12,14). Although EA is reasonably well-tolerated in NHPs, the Laflamme group (14) reported that EA can be lethal in pigs. For this reason, EA has emerged as the biggest impediment to the clinical translation of human cardiomyocyte transplantation (15).

We hypothesized that the risk of EA may be mitigated by treatment with clinically available anti-arrhythmic drugs. Because the pig shows heightened sensitivity to EAs, is a well-established model in cardiovascular research (16) and cell therapy (17), whose larger heart permits the use of percutaneous delivery catheters, we chose to test the hypothesis in this large animal model. In the first phase of our study, we screened a panel of seven anti-arrhythmic agents. The broad-acting (class III) antiarrhythmic amiodarone and the pacemaker inhibitor (class 0) ivabradine emerged independently as the most promising agents for control of rhythm and rate, respectively. We therefore performed a second phase to test the effect of combined amiodarone and ivabradine treatment. We found that this regimen reduced sudden cardiac death, as well as suppressed tachycardia and arrhythmia.

## Methods

### hESC-CM production

These studies were approved by the University of Washington Stem Cell Research Oversight Committee. Two lines of hESCs were used in this study. Initial subjects received H7 (WiCell)-derived cardiomyocytes that were cultured, expanded, and differentiated in suspension-culture format by collaborators at the Center for Applied Technology Development at the City of Hope in California, all as previously described (11,12,18). Most subjects received RUES2 (Rockefeller University)-derived cardiomyocytes produced in our laboratory in stirred suspension culture format. Briefly, RUES2 hESC were cultured to form aggregates and were expanded in commercially available media (Essential 8, Gibco). For cardiac differentiation, suspension adapted pluripotent aggregates were induced to differentiate in RPMI-1640, MCDB-131, or M199 supplemented with B-27 (all from Gibco) or serum albumin, by timed use of small molecule GSK 3 inhibitors and Wnt/β-catenin signal pathway inhibitors (Tocris). Twenty-four hours prior to cryopreservation, RUES2 hESC-CMs were heat-shocked to enhance their survival after harvest, cryopreservation, thaw, and transplantation. Cardiomyocyte aggregates were dissociated by treatment with Liberase TH (Fisher) and TrypLE (Gibco) and were cryopreserved in CryoStor CS10 (Stem Cell Technologies) supplemented with 10 μM Y-27632 (Stem Cell Technologies) using a controlled-rate liquid nitrogen freezer. Approximately 3 h before transplantation, cryopreserved hESC-CM were removed from cryogenic storage (−150 °C to −196 °C) and thawed in a 37 °C water bath (2 min ± 30 s). RPMI-1640 supplemented with B-27 and ≥200 Kunitz Units/mL DNase I (Millipore) was added to the cell suspension to dilute the cryopreservation media.

Subsequent wash steps were done using RPMI-1640 basal media in progressively smaller volumes to concentrate the cell suspension. For the last centrifugation step, the cell pellet was resuspended in a sufficient volume of RPMI-1640 to achieve a target cell density for injection of ~3 ×10^9^ cells/mL in 1.6 mL. The final volume of the cell suspension was determined by the results of a count sampled before the final centrifugation step. Cell counts were performed as described previously to achieve a final total dose of 500×10^6^ live cells per transplant (12).

### Study design

The objective of this study was to identify a pharmacological regimen to attenuate arrhythmias following cardiac remuscularization therapy. This study was designed in two phases: the first to observe the natural history of EA in the minipig model and screen various antiarrhythmic agents for possible efficacy, and the second to rigorously test for efficacy of selected candidates (Fig. 1). All subjects were 30–40 kg castrated male Yucatan minipigs between 6–13 months of age (Premier BioSource). In Phase 1, nine subjects underwent cardiac remuscularization therapy with 500 × 10^6^ hESC-CMs delivered by direct surgical trans-epicardial injections or, later by percutaneous trans-endocardial injections (Table 1).

**Fig. 1.**
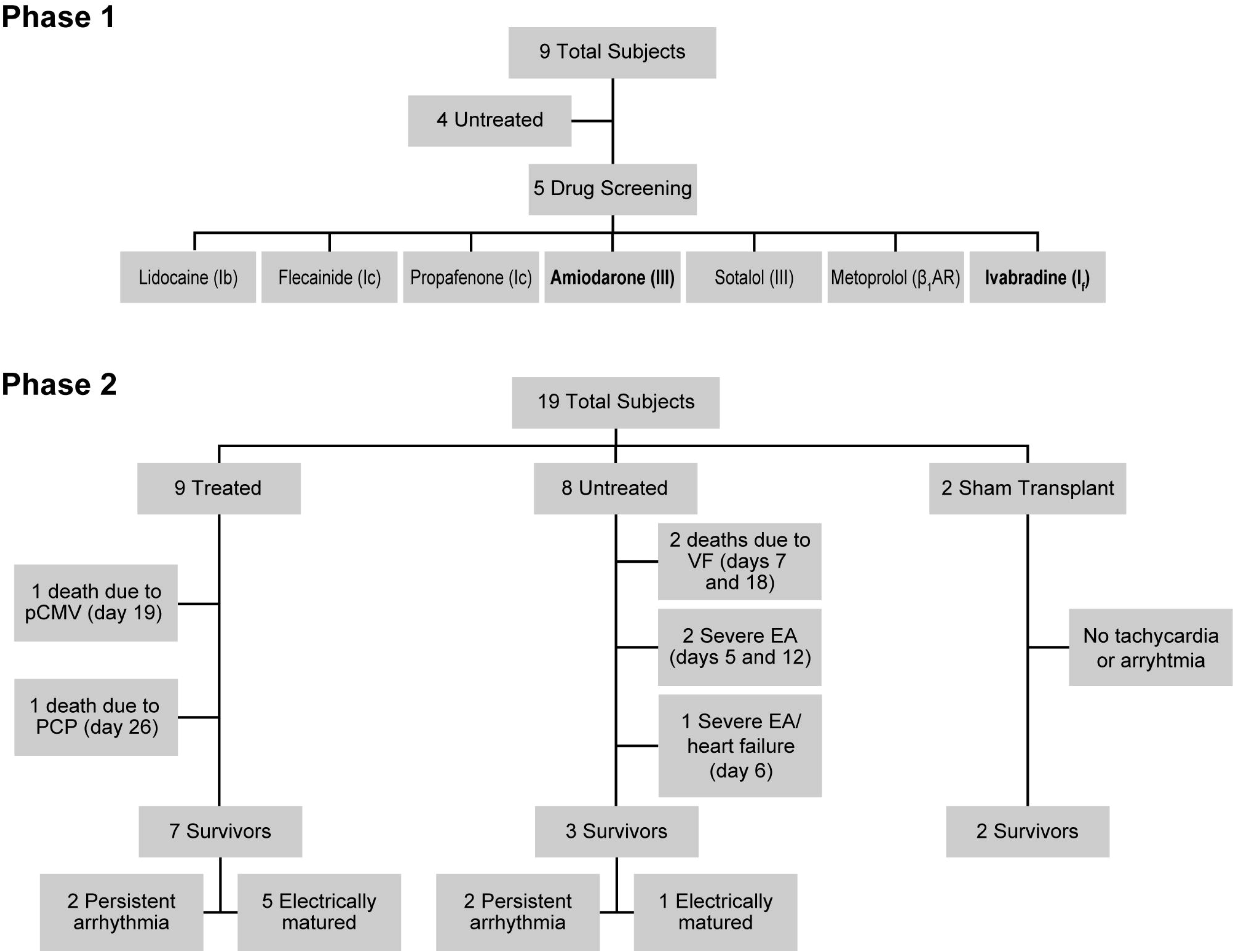
Study Design Flowchart of the study design. Phase 1 consisted of nine total subjects, four untreated with any antiarrhythmic to study the natural history of engraftment arrythmia (EA) and five used to screen seven candidate antiarrhythmic agents. Amiodarone and ivabradine were found to have promising signals of effect and advanced for further study. Phase 2 consisted of 19 total subjects: nine treated with amiodarone and ivabradine and eight untreated following hESC-CM transplantation, and two untreated following sham transplantation.

**Table 1.**
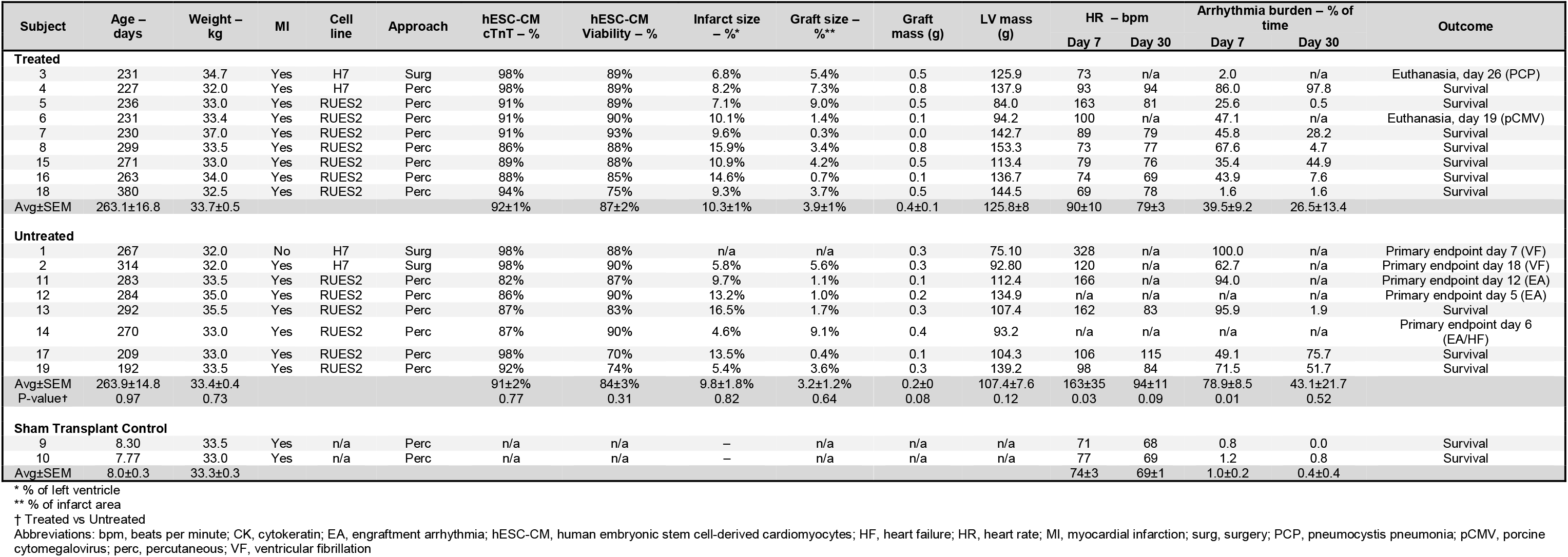
Characteristics of Study Subjects and Clinical History.

The first four subjects (one non-infarcted and three infarcted) were followed to learn the natural history of EA and establish clinical endpoints and parameters for the Phase 2 drug trial. The subsequent five subjects underwent systematic dosing with antiarrhythmic agents with continuous electrocardiography (ECG) monitoring to determine effect on rhythm and rate (Online Table 1, Online Methods). Among the nine subjects in Phase 1, we observed high mortality, with six out of nine experiencing ventricular fibrillation (VF) or tachycardia-induced heart failure requiring euthanasia. VF typically followed frequent episodes of unstable EA >350 beats per minute (bpm), and tachycardia-induced heart failure requiring euthanasia was characterized by chronically elevated heart rates >150 bpm.

In Phase 2, we conducted a two-drug antiarrhythmic study with amiodarone and ivabradine, enrolling an additional 17 subjects (9 treated, 8 untreated) that underwent MI and percutaneous transplantation with hESC-CMs at two weeks post-MI. Two additional subjects underwent MI with sham vehicle injection to serve as sham transplant controls (Fig. 1, Online Fig. 1). The primary endpoint was prespecified as combined cardiac death (either spontaneous death from arrhythmia or heart failure, or clinically directed euthanasia necessitated by sustained tachycardia >350 bpm or signs of heart failure). Prespecified secondary endpoints were suppression of tachycardia, percent time in arrhythmia (arrhythmia burden) and resolution of arrhythmia, termed “electrical maturation” and defined as arrhythmia burden <25% for 48 consecutive hours. Antiarrhythmic therapy was discontinued after electrical maturation or post-transplantation day 30, whichever was earlier. To prevent tachycardia-induced cardiomyopathy, we titrated ivabradine treatment to maintain target heart rate <150 bpm. Based on early experience that tachycardia >350 bpm often degenerated to VF, subjects were euthanized humanely if heart rates >350 bpm were reached. Continuous telemetric ECG was monitored for eight weeks total (two weeks post-MI and six weeks post-transplantation). Of note, subjects 1 and 2 (untreated) and 3 and 4 (treated) received H7 hESC-CM and subjects 1, 2 and 3 were transplanted surgically prior to adopting percutaneous delivery. Subject 5 was euthanized on day 37 as a prespecified endpoint following electrical maturation, prior to extending the study duration to 6 weeks post-transplantation for extended treatment washout and monitoring.

### Cardiac remuscularization therapy

Cell transplantation for our three initial subjects (1–3) was performed by direct transepicardial injection into the peri-infarction region as previously described for NHP with minor modification (12). Briefly, a partial median sternotomy was performed to expose the infarcted anterior left ventricle. Purse-string sutures were preplaced at five discrete locations subtended by the LAD, targeting the central ischemic region and lateral border zones. After cinching the purse-string tightly around the needle, three injections of 100 μL each were performed by partial withdrawal and lateral repositioning, for a total of 15 injections to deliver total dose of 500 × 10^6^ hESC-CMs. All subsequent subjects (4–19) received cell transplantation via percutaneous trans-endocardial injection using the NOGA-MyoStar platform (BioSense Webster) to first map the infarct region in the left ventricle, and then to deliver 16 discrete endocardial injections of 100 μL each for total dose of 500 × 10^6^ hESC-CMs. Injections were only performed with excellent location and loop stability, ST-segment elevation and presence of premature ventricular contraction (PVC) with needle insertion in an appropriate location by electroanatomical map and unipolar volage. For both surgical and percutaneous cell transplantation, two-thirds of injections were placed into the peri-infarct border zone defined by unipolar voltage of 5–7.5 mV and the remaining one-third into the central ischemic region defined as unipolar voltage of < 5 mV. Two subjects (9 and 10) were infarcted as per protocol but received sham injections of RPMI-1640 vehicle without cells to serve as sham transplant controls.

### Antiarrhythmic treatment

The treated cohort was loaded with oral amiodarone 1000–1200 mg orally twice daily starting seven days prior to cell transplantation followed by maintenance dose of 400–1000 mg orally twice daily to maintain a steady-state plasma level of 1.5–4.0 μg/mL (Online Fig. 2). Ivabradine was started at 2.5 mg orally twice daily when sustained tachycardia reached ≥ 150 bpm and titrated every 3 days up to 15 mg twice daily for goal heart rate < 125 bpm. All but one subject (1) required adjunctive ivabradine for additional heart rate control. Antiarrhythmics were discontinued after electrical maturation was achieved or post-transplantation day 30, whichever was earlier, to allow for treatment washout and assess for recrudescence of arrhythmia. All subjects tolerated the antiarrhythmic regimen without complication. Untreated and sham transplant control subjects did not receive antiarrhythmic agents following the MI procedure, but otherwise received all immunosuppression and standard care.

**Fig. 2.**
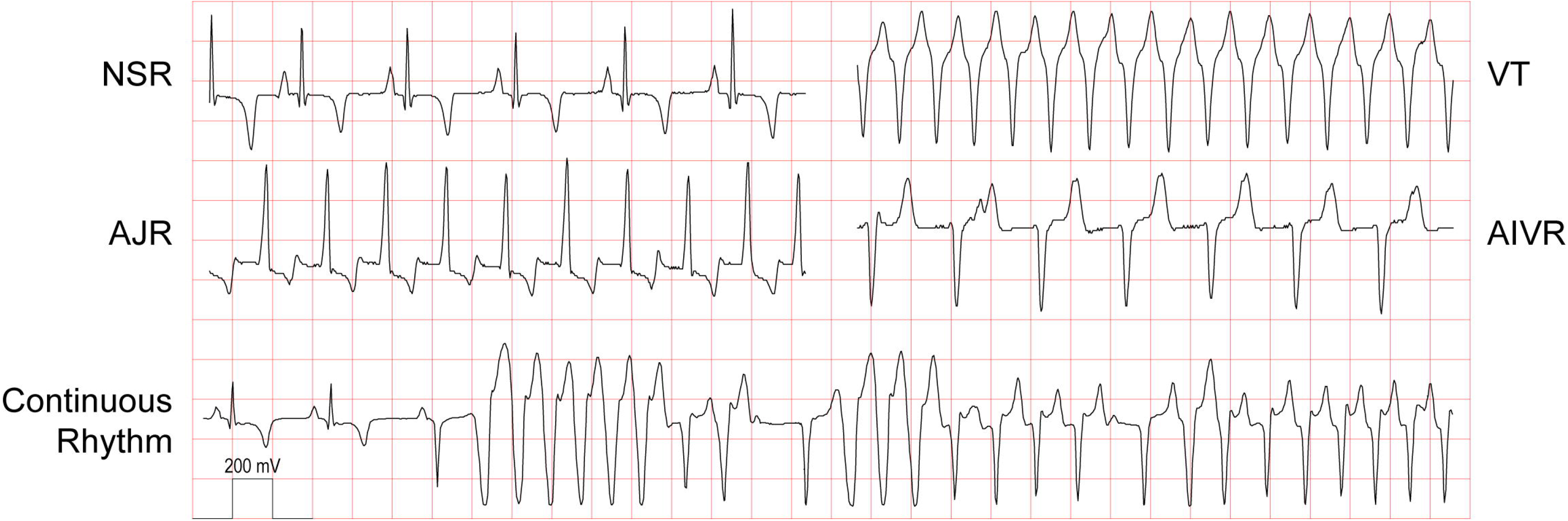
Engraftment Arrhythmia Variable morphologies of engraftment arrhythmia (EA) in a single minipig. Examples of normal sinus rhythm (NSR) and three morphologies of EA resembling accelerated junctional rhythm (AJR), ventricular tachycardia (VT) and accelerated idioventricular rhythm (AIVR) are observed in a single untreated subject (12). Note the variation in rate, electrical axis, and QRS duration. A continuous rhythm recording below exhibits polymorphic EA with QRS complexes varying in rate, duration, and electrical axis. No sustained arrythmias were noted in surgical sham controls. 1 box vertical 200 mV, horizontal 0.2 sec.

### Statistical analysis

Statistical analyses and graphing were performed using Prism 8.4.2 software (GraphPad) and Stata 15 (StataCorp, College Station, Texas). Data are presented as mean ± standard error of the mean (SEM). Comparisons were performed using Mann-Whitney test with significance threshold of P < 0.05 unless otherwise specified. The sample size to demonstrate a difference in mortality rate of 67% (untreated group) versus 0% (treatment group), with alpha 0.05 and 90% power, was estimated to be 8 per cohort. Kaplan-Meier plots show survival curves for the primary endpoint of cardiac death, unstable EA or heart failure, and for all-cause mortality. Cox proportional regression models are used to estimate the hazard ratio (HR) between the two treatment groups, for the primary outcome and for mortality. Significance is based on the likelihood ratio test and confidence intervals on HR are computed by inverting the likelihood test, based on varying the offset term in the stcox procedure in Stata.

## Results

### Clinical history of engraftment arrhythmia

A flow chart for all subjects in the study is shown in Fig. 1 and clinical summaries are provided in Table 1. No significant arrhythmias were noted in the two untreated sham transplant control subjects (9 and 10) that underwent myocardial infarction and percutaneous intracardiac injection of vehicle. All subjects that received human cardiomyocyte grafts developed EA between 2–6 days following cell transplantation. Initiation of EA was characterized by salvos of non-sustained VT, and this typically progressed to periods of sustained VT with rates ranging from 110 to 250 bpm (Fig. 2). The VT was often polymorphic, with the same subject showing different electrical axes and both wide- and narrow-complex tachycardia at different times. In four of the eight untreated subjects, EA was either fatal or necessitated euthanasia due to a prespecified endpoint of unstable tachycardia (defined as sustained heart rate > 350 bpm). In one additional untreated case (subject 12), acute heart failure was noted clinically shortly after initiation of EA at a rate of 300 bpm, and based on recommendations from veterinary staff, the subject was euthanized. Signs of heart failure were subsequently confirmed on necropsy. In all other cases, EA was noted with a rapid acceleration to > 350 bpm (subjects 11 and 12) and, in two cases, deterioration to VF prior to euthanasia (subjects 1 and 2) (Table 1). Three out of four arrhythmic endpoints occurred within the first three days of developing EA, and they occurred when tachyarrhythmia was nearly constant. Mean heart rate peaked at day 8 post-transplantation and began to decline after that, whereas the arrhythmia burden plateaued from days 8–16 and began to normalize thereafter. Of the three survivors in the untreated cohort, two did not normalize rhythm and experienced on average 42% arrhythmia burden at the end of study (subjects 15 and 17). The single subject in the untreated cohort that normalized heart rate and rhythm did so on day post-transplant day 26 (subject 11).

### Screening drugs for anti-arrhythmic effects

In Phase 1 of the study, we screened six canonical antiarrhythmic agents broadly targeting sodium channels, potassium channels, and beta-adrenergic receptors: lidocaine (class Ib Vaughan-Williams-Singh antiarrhythmic, sodium channel inhibitor), flecainide (Ic, sodium channel inhibitor), propafenone (Ic, sodium channel inhibitor), amiodarone (III, potassium channel inhibitor), sotalol (III, potassium channel inhibitor) and metoprolol (β1-adrenergic receptor inhibitor) for effect on EA heart rate and rhythm. In addition, the funny current/HCN4 channel antagonist, ivabradine, was tested (Online Table 1). This series was not meant to be comprehensive but rather to rapidly identify candidate agents.

Animals were brought into the laboratory while in EA, anesthetized, and the effects of short-term intravenous infusion or oral treatment of anti-arrhythmic agents were studied. In three instances, intravenous amiodarone successfully cardioverted unstable EA from >350 bpm to a lower heart rate, typically including brief episodes of sinus rhythm (Fig. 3A). Oral ivabradine demonstrated robust dose-dependent effects on heart rate but did not restore sinus rhythm (Fig. 3B). Four of the other drugs had no significant effect in this screen (lidocaine, flecainide, sotalol, and metoprolol). Propafenone briefly reduced heart rate and restored sinus rhythm in two drug challenges, but this drug was associated with substantial gastrointestinal toxicity and not studied further (data not shown).

**Fig. 3.**
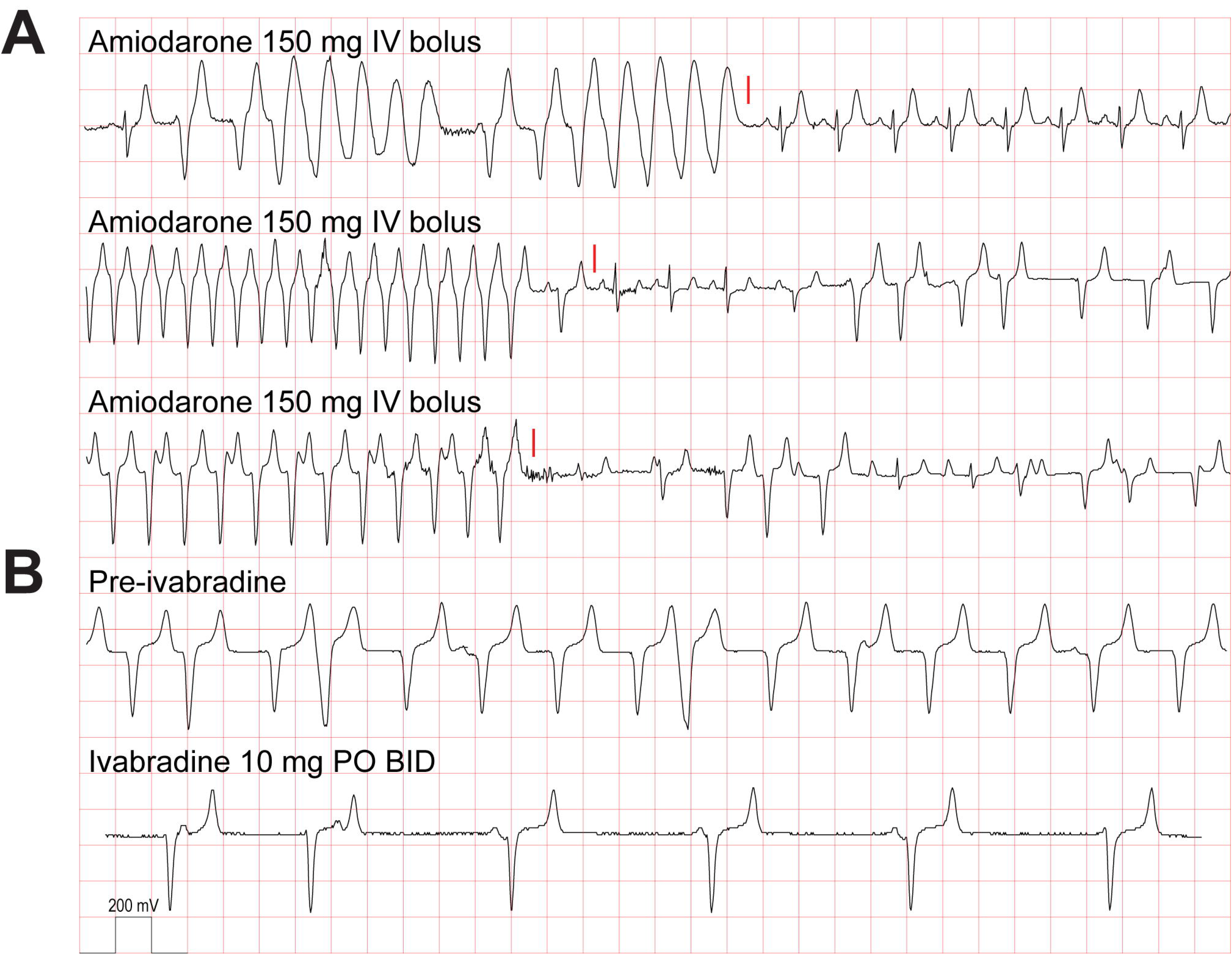
Acute effects of amiodarone and ivabradine Acute effects of amiodarone and ivabradine on engraftment arrhythmia. Amiodarone was effective as an intravenous bolus to cardiovert engraftment arrythmia to normal sinus or a lower heart rate transiently in three separate instances (A, red line). Ivabradine administered orally significantly slowed EA but did not cardiovert two days following initiation (B). These data supported a combined amiodarone and ivabradine antiarrhythmic strategy for rhythm and rate control of EA. 1 box vertical 200 mV, horizontal 0.2 sec.

### Amiodarone-ivabradine enhance survival

Given their distinct mechanisms of action and complementary effects on heart rate and rhythm, we formally tested the hypothesis that amiodarone along with ivabradine would reduce a combined primary endpoint of cardiac death, unstable EA >350 bpm and heart failure in Phase 2 of the study. A total of nine treated, eight untreated, and two sham transplant subjects were enrolled in the study with similar baseline and cell transplantation characteristics (Table 1). As detailed in the Methods, treated animals received bolus and maintenance doses of amiodarone, and ivabradine was given as needed to keep heart rates <150 bpm. All treated subjects survived without the primary cardiac endpoint compared to 3/8 (37.5%) of untreated subjects (Fig. 4A). The hazard ratio of the primary endpoint was 0.000 (95% CI, 0.000–0.297; p = 0.002) with antiarrhythmic treatment. Of note, two of the treated subjects (3 and 6) experienced non-cardiac deaths at post-transplant days 19 and 26 due to immunosuppression-related complications (Pneumocystis pneumonia and porcine cytomegalovirus, respectively). Intention-to-treat analysis of overall survival also favored the treated cohort with hazard ratio of 0.212 (95% CI, 0.030– 1.007; p = 0.05) (Fig. 4B).

**Fig. 4.**
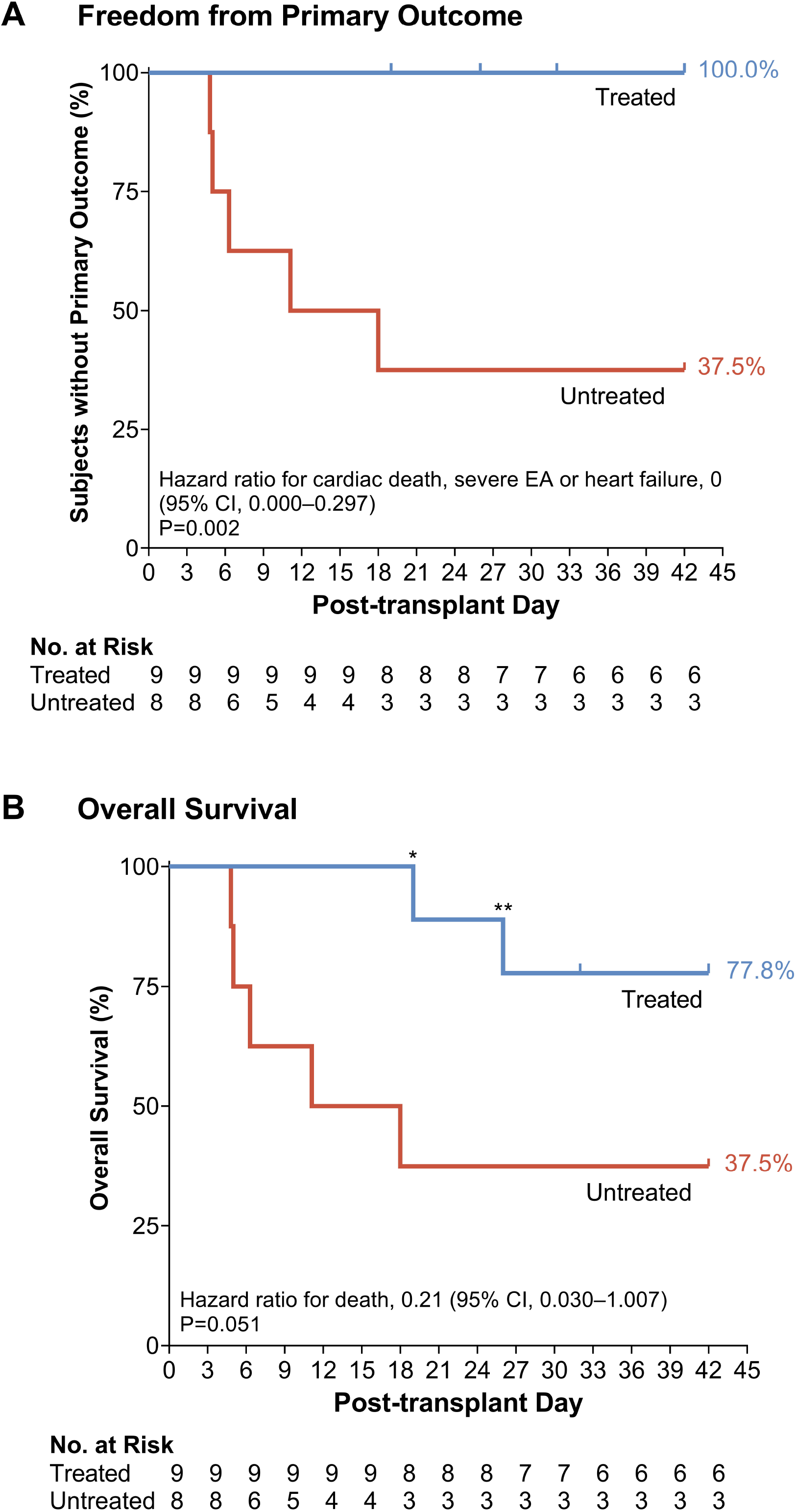
Kaplan–Meier Survival Curves Antiarrhythmic treatment with amiodarone and ivabradine for engraftment arrhythmia in pig. (A) Kaplan-Meier curve for freedom from primary outcome of cardiac death, unstable EA or heart failure was significantly improved in the treated compared to untreated cohort (p=0.002). Tic marks on treatment line indicate non-cardiac death due opportunistic infection (days 19 and 26) or a planned euthanasia (day 30). (B) Kaplan-Meier curve for overall survival shows statistically borderline improvement in the treated compared to untreated cohort (p=0.051). *Death due to Pneumocystis pneumonia. **Death due to porcine cytomegalovirus. Abbreviations: CI, 95% confidence interval.

### Suppression of tachycardia and arrhythmia burden

Pooled and individual subject-level data of heart rate and arrhythmia burden are provided in Figs 5A/5B and 5C/5D, respectively. The average heart rate was significantly lower with antiarrhythmic treatment compared to no treatment. Mean heart rates peaked at post-transplantation day 7 in untreated animals at 163 ± 35 bpm, versus average heart rates of 90 ± 10 bpm in the treated cohort (p = 0.03) (Table 1 and Fig. 5). Heart rate in the treated animals was not significantly different than the normal resting heart rate prior to MI and transplant (84 ± 1 bpm, p = 0.21). Following transplantation, peak daily heart rate for the study duration averaged 305 ± 29 bpm in untreated animals, whereas treatment significantly attenuated peak heart rate to 185 ± 9 bpm (p = 0.005) (Fig. 5E). We defined arrhythmia burden as the percentage of the day spent in arrhythmia. Treatment reduced peak arrhythmia burden from 96.8 ± 2.9% to 76.5 ± 7.9% (p = 0.003) (Fig. 5F). No differences in heart rate or arrhythmia burden were noted at post-transplant day 30, as the majority of arrhythmia had resolved irrespective of treatment (Fig. 5A/B) (p = 0.09 and p = 0.52, respectively).

**Fig. 5.**
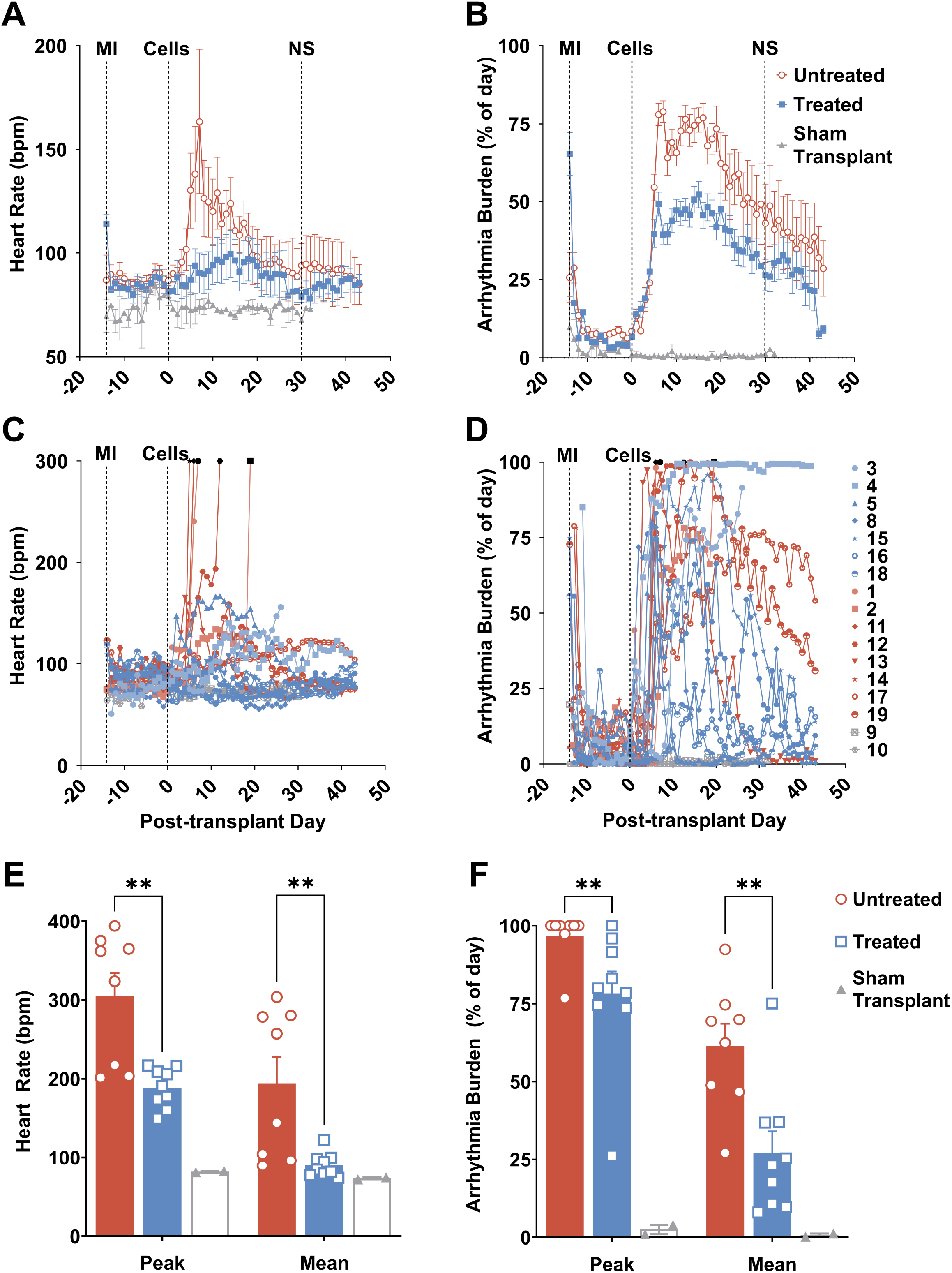
Heart Rate and Engraftment Arrhythmia Burden Effect of antiarrhythmic treatment on heart rate and arrhythmia burden. Pooled daily average heart rate (A) and pooled daily average arrhythmia burden (B) in treated (blue) compared to untreated (red) cohort. The difference in heart rate or arrhythmia burden between treated and untreated cohorts was not significant (NS) by day 30 post-transplantation. Sham transplant (grey) did not induce tachycardia or arrythmia. Subject-level averaged daily heart rate (C) and arrhythmia burden (D) for antiarrhythmic treated (blue), untreated (red) and sham transplant (grey). Unexpected death or euthanasia denoted by black symbol. Overall peak and mean daily heart rate (E) and overall peak and mean daily arrythmia burden (F) were significantly reduced in treated (blue) compared to untreated (red) cohorts. No tachycardia or arrhythmias were noted in the sham transplant control cohort. ** p ≤ 0.01.

Antiarrhythmic treatment was safely discontinued between day 24–34 in all treated subjects that achieved electrical maturation without recrudescence of arrhythmia (Fig. 5). Two treated and two untreated subjects (3, 4 and 15, 17, respectively) failed to mature electrically and exhibited significant arrhythmia at the end of study. In these four animals, heart rates were well controlled irrespective of treatment, and they survived until the study’s completion. Average serum amiodarone was sub-therapeutic at 0.42 ± 0.12 μg/mL within 1 week of discontinuation (Online Fig. 2).

### Percutaneous delivery of hESC-CMs in infarcted porcine model

Catheter-based endocardial delivery of hESC-CMs was safe and effective in remuscularizing the infarcted porcine heart (Online Fig. 3). No significant differences in myocardial infarct or cardiomyocyte graft sizes were observed between the treatment groups. The average infarct size for the treated and untreated cohorts were comparable at 11.7±1.1% and 10.5±2.0% of the left ventricle, respectively (p=0.59). Graft size relative to infarct size was also comparable at 2.3±0.7% and 2.8±1.3% for treated and untreated, respectively (p= 0.74). Delivery of hESC-CM successfully targeted the peri-infarct border zone and central ischemic regions as intended and resulted in discrete hPSC-CM grafts transplanted into host myocardium, as previously reported (11–14). All grafts localized to the anterior, antero-septal and antero-lateral walls and as previously reported in pig (14), appeared structurally immature at early time points before two weeks post-transplantation with increasing maturity up to the end of study.

### Graft interaction with host Purkinje conduction system

The narrow-complex tachycardia that resembles accelerated junctional rhythm (Fig. 2) was not observed in our previous NHP studies (11,12) but was common in the minipig. Pigs are known to have an extensive Purkinje fiber network that extends transmurally throughout the ventricular myocardium, whereas in macaques and humans the Purkinje network is subendocardial (19,20). We hypothesized that the narrow-complex VT resulted from graft automaticity conducting through intramural Purkinje fibers and propagating to the rest of the ventricle. Histology confirmed the mesh-like network of intramural Purkinje fibers (PFs) throughout the minipig left ventricle (Online Fig. 4A, Online Video 1). There were multiple examples of hESC-CM grafts in direct contact with these intramural branches of the Purkinje system. (Fig. 6, Online Video 2). We used connexin 40 (Cx40) immunostaining to specifically stain Purkinje fiber gap junctions (19,21), and confirmed their identity by their reduced myofibril content and the absence of T tubules (Online Fig. 4B). This supports the hypothesis that the pig’s unique Purkinje network anatomy contributes to narrow-complex engraftment arrhythmia.

**Fig. 6.**
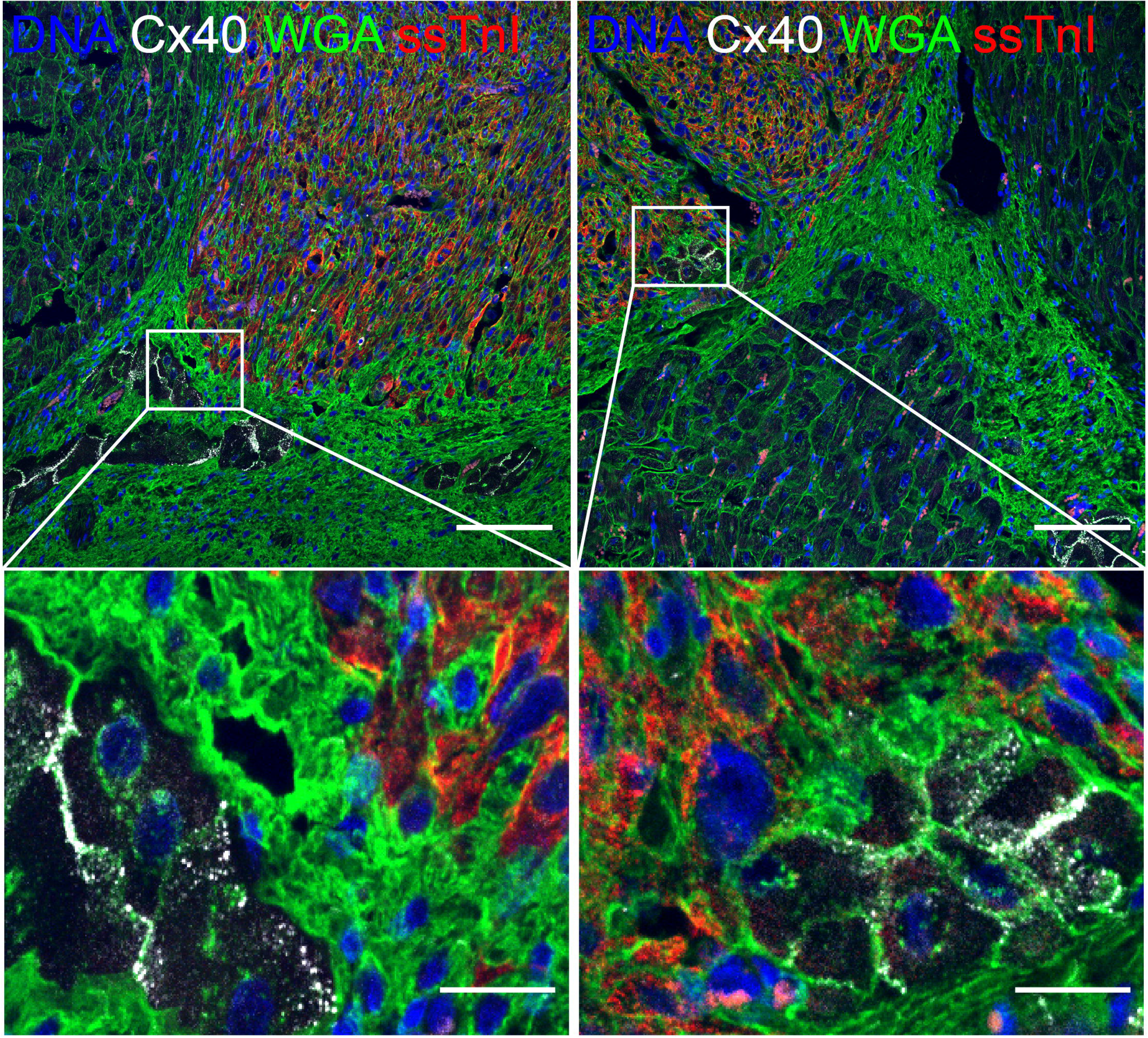
Graft interaction with host Purkinje conduction system Transplanted hESC-CM grafts interact with a diffuse Purkinje conduction system in the porcine myocardium. hPSC-cardiomyocyte graft 2-weeks post-transplantation marked by human-specific slow skeletal cardiac troponin I (ssTnI, red) interact with Cx40-positive (white) Purkinje fibers. Purkinje-transitional cell-graft (left panel) and direct Purkinje-graft (right panel) interaction are observed, scale bar 100 μm or 20 μm (magnified).

## Discussion

Intramyocardial transplantation of hPSC-CM is a promising strategy to remuscularize the infarcted heart and restore function (5). Such a therapy to prevent and treat heart failure would be a seminal advance in addressing a large unmet clinical need. Studies in large animals have demonstrated long-term efficacy but also defined a significant safety signal of generally transient but potentially fatal arrhythmias. As demonstrated in earlier studies (12–14), EA is a predictable complication of cardiac remuscularization therapy for myocardial infarction (22). In the NHP, EA typically presents as a wide-complex tachycardia with a variable electrical axis (11,12), and this was reproduced in the minipig recently by Laflamme laboratory (14). Here, we further describe EA as polymorphic and interpret the changes in electrical axis as ectopy originating from different graft foci. Interestingly, in the pig we also observed a narrow-complex VT that alternated with a wide-complex tachycardia, a pattern not seen in the NHP. Histology of native and grafted porcine myocardium support the hypothesis that the wide-complex beats originate from grafts interacting with the working cardiac myocytes with slow conduction, and that the narrow complex beats originate when grafts interact with the intramural Purkinje fibers that diffusely permeate the porcine heart (19,20).

All 17 subjects transplanted with 500 × 10^6^ hESC-CMs demonstrated significant burden of arrhythmia that, while typically transient, was associated with high mortality in pigs. We observed higher morbidity and mortality related to EA than the recent study by Laflamme and colleagues (14), perhaps reflecting differences in our animal model including use of Yucatan minipigs, percutaneous cell delivery, or our cell product. Our experience with this model suggests two primary mechanisms of cardiac morbidity. Firstly, rapid EA > 350 bpm often degenerates to fatal ventricular fibrillation, and secondly, heart failure commonly ensues in pigs with chronic tachycardia > 230 bpm (23). Consequently, our primary endpoint included these parameters to limit excessive mortality in our antiarrhythmic trial.

Combined antiarrhythmic treatment with baseline amiodarone and adjunctive ivabradine safely prevented the combined primary endpoint of cardiac death, unstable EA and heart failure in all treated subjects, indicating that the risk of EA may be mitigated through pharmacology. Treatment was associated with significantly decreased peak tachycardia and arrhythmia. Once subjects experienced sustained improvement in arrhythmia burden, termed electrical maturation, antiarrhythmic therapy was successfully withdrawn in all subjects. Thus, short-term amiodarone and ivabradine treatment promoted electrical stability until the grafts became less arrhythmogenic.

The mechanism of benefit for our antiarrhythmic treatment may be related to suppression of automaticity, reducing both heart rate and arrhythmia burden. The drugs were particularly beneficial during the early phase of EA, which carries the greatest risk of deterioration to VF. Electrophysiological studies performed by our group and the Laflamme laboratory in NHP (12) and pig (14), respectively, suggests that the etiology of EA is increased focal automaticity, rather than macro-reentry typically observed with clinical ventricular tachycardia (24). As EA became unstable in the untreated animals, heart rates rapidly accelerate to >350 bpm, and we cannot exclude the possibility that this escalation could have a distinct mechanism, e.g., automaticity leading to reentry. This may explain why treatment successfully suppressed unstable and fatal arrhythmias but was unable to prevent EA altogether.

The efficacy of ivabradine to rate-control EA suggests that its pharmacologic target, the I_f_ current carried by the HCN4 channel, which is highly expressed in immature cardiomyocytes and hPSC-CMs (25), may be an important mediator. Ivabradine, by itself, never abrogated EA, suggesting that while the I_f_ current can accelerate the rate of EAs, I_f_ is not the current responsible for generating the ventricular automaticity triggered by engraftment. In contrast, amiodarone reduced the burden of EA chronically and clearly restored sinus rhythm in some acute infusion experiments (Fig. 3). Although classified principally as a K^+^ channel blocker (class III), amiodarone is well-known also to antagonize Na^+^ channels, Ca^2+^ channels, and β-adrenergic receptors (26). Thus, it is difficult to gain insights into the mechanism of EA from amiodarone’s efficacy. The disappearance of EA coincides with maturation of the stem cell-derived graft (11,12,27), and we with others have hypothesized that the window of arrhythmogenicity may reflect a period of *in vivo* graft maturation prior to reaching a state more similar to host myocardium (25,28–32). Additional strategies such as promoting maturation prior to transplantation, gene editing, and modulating host/cell interaction may provide additional means of arrhythmia control and a comprehensive protocol invoking multiple complementary mechanisms of action may ultimately be necessary to ensure safety. Further investigation of the etiology of EA would be accelerated by the development of higher throughput *in vivo*, *ex vivo*, *in vitro* and/or *in silico* platforms to perform genetic, pharmacological, electrophysiological and/or modeling studies before phenotyping in large animal models.

Engraftment arrhythmia is the most significant barrier to clinical translation of cardiac remuscularization therapy. The natural history of EA emerging from the NHP and more recent porcine data suggests that, once EA resolves, there is low risk for further arrhythmia. This study provides a proof-of-concept that clinically relevant antiarrhythmic drug treatment can successfully suppress fatal arrhythmias and control tachycardia to achieve electrical quiescence. This could be an important tool toward reaching an acceptable safety profile for clinical development.

While this study demonstrates that EA is responsive to pharmacologic suppression, there are several limitations. It would be useful to perform a longer follow-up to establish the long-term effectiveness of EA mitigation as well as dosing studies to optimize the treatment regimen. We did not randomize enrollment of animals or assess whether sex is a biological variable. Although we took pains to administer clinically relevant doses of amiodarone and ivabradine, we cannot exclude the possibility that EA in itself is dependent on the dose of cells transplanted. The dose utilized is comparable to that utilized to demonstrate long-term function benefit in NHP (500 × 10^6^ versus 750 × 10^6^ hESC-CMs, respectively),(12) but dosing studies have not been reported. Future studies will also ideally include functional endpoints to determine mechanical efficacy with background guideline-directed medical therapy such as inhibitors of the renin–angiotensin–aldosterone and β-adrenergic systems.

## Conclusions

In this study utilizing a porcine infarction model of cardiac remuscularization therapy, EA was universally observed and associated with significant mortality. Chronic amiodarone treatment combined with adjunctive ivabradine successfully prevented the combined primary endpoint of cardiac death, unstable EA and heart failure. Overall survival was significantly improved with antiarrhythmic treatment and associated with heart rate and rhythm control. The mechanisms of engraftment arrhythmia remain incompletely understood and merit concerted scientific inquiry.

## Abbreviations

CI: Confidence interval
EA: Engraftment arrhythmia
ECG: Electrocardiogram
hESC-CM: Human embryonic stem cell-derived cardiomyocyte
hPSC-CM: Human pluripotent stem cell-derived cardiomyocyte
MI: Myocardial infarction
NHP: Non-human primate
PF: Purkinje fiber
VF: Ventricular fibrillation

## Author contributions

K.N. conceived the study, led experimental design, performed surgical and percutaneous procedures, analyzed and interpreted data, and wrote the manuscript. L.E.N. assisted the conception, experimental design and execution of the study, performed surgical and percutaneous procedures, supervised veterinary care, contributed to data analysis and edited the manuscript. X.Y. performed histologic analysis and edited the manuscript. G.J.W. performed histologic analysis and contributed to the manuscript. D.E performed histologic analysis and contributed to the manuscript. H.T. assisted surgical and percutaneous procedures, assisted experimental design and performed histologic analysis. S.D. assisted surgical and percutaneous procedures and performed histologic analysis. F.A.K. cultured and characterized the hESC-CM cell product. A.J. cultured and characterized the hESC-CM cell product. A.F. cultured and characterized the hESC-CM cell product. D.S.N. assisted experimental design and writing the manuscript. S.M. assisted experimental design and edited the manuscript. A.B. assisted experimental design and edited the manuscript. M.R.R. assisted the study’s conception, experimental design and execution. K.C. performed statistical analysis and contributed to the manuscript. H.R. assisted the study’s conception, experimental design and execution. L.P. assisted the study’s conception, experimental design and execution. B.C.K. assisted the study’s conception and experimental design. S.K. assisted the study’s experimental design and execution and supervised cell manufacturing. R.S.T. assisted the study’s conception, experimental design and execution, supervised cell manufacturing and obtained research funding. W.R.M. assisted the study’s conception, experimental design and execution and obtained research funding. C.E.M. conceived and supervised the study, obtained research funding and contributed to data analysis and writing the manuscript.

## Acknowledgements

These studies were supported by the UW Medicine Heart Regeneration Program, the Washington Research Foundation, and a gift from Mike and Lynn Garvey (all Seattle, WA). This work also was supported in part by NIH Grants R01HL128362, and a grant from the Fondation Leducq Transatlantic Network of Excellence (Boston, MA; to C.E.M.) and the Bruce-Laughlin Research Fellowship (Seattle, WA; to K.N.). We are indebted to Emily Spaulding, Gary Fye, Piper M. Treuting, Thea Brabb and colleagues in the UW Department of Comparative Medicine for their exceptional veterinary care and consultation. We acknowledge the Center for Applied Technology Development at the City of Hope (Duarte, CA) for providing H7 hESC-CM used for our studies and use of these cells was funded in part through the National Heart Lung and Blood Institute’s Production Assistance for Cell Therapies (PACT) (Bethesda, MD). We thank the Garvey Imaging Core at UW for assistance with microscopy.

## Central Illustration

Fig. 4.

## Clinical Perspectives

Heart failure remains a significant cause of morbidity and mortality following myocardial infarction (MI). Cardiac remuscularization with transplantation of pluripotent stem cell-derived cardiomyocytes is a promising preclinical therapy to restore function. Recent large animal data, however, have revealed a significant risk of engraftment arrhythmia (EA). The present study provides proof-of-concept evidence that a combination of amiodarone and ivabradine can effectively prevent EA-related mortality and suppresses tachycardia and arrhythmia burden. Thus, pharmacologic suppression of EA may be a viable strategy to improve safety and allow further clinical development of cardiac remuscularization therapy.

**Online Fig. 1 Study timeline Study timeline for Phase 2 drug trial of chronic amiodarone and adjunctive ivabradine therapy.** Myocardial infarction (MI) was induced by 90-minute balloon occlusion of the mid-left anterior descending artery two weeks prior to human embryonic stem cell-derived cardiomyocyte transplantation (day 0). All subjects received multi-drug immunosuppression. Treated cohort received rate and rhythm control with combined oral amiodarone and adjunctive oral ivabradine.

**Online Fig. 2 Amiodarone Levels Plasma amiodarone levels in pigs.** Amiodarone levels were measured in plasma by a custom liquid chromatography-mass spectrometry assay. Chronic oral amiodarone in six pigs was discontinued after achieving electrical maturation and stabilization of engraftment arrythmia. Serum through concentrations of amiodarone were assayed weekly including 3–4 weeks after discontinuation.

**Online Fig. 3 Graft Histology and Injection Location hESC-CM graft histology and location.** Left panel: Histological sections stained with picrosirius red to identify collagen (infarct) and fast green to identify viable myocardium. Adjacent sections labeled with human cTnT (brown) identify transplanted hESC-CM graft within unstained porcine myocardium and scar tissue. Sections from both treated and untreated subjects were obtained on post-transplantation day 42. Right panel: Transplanted hESC-CM grafts were located similarly between treated (blue) and untreated (red) cohorts and successfully targeted the infarct and peri-infarct regions of the anterior wall.

**Online Fig. 4 Purkinje fiber staining in native pig myocardium Purkinje fibers are distributed in a mesh-like network throughout the native porcine myocardium and are specifically marked by Connexin 40.** Subendocardial and intramyocardial connexin 40 (Cx40)-positive Purkinje fibers (PFs, white) in transverse section of left ventricular free wall, scale bar 2 mm (A). Intramyocardial PFs are shown with higher magnification insets. Further magnified view of white boxed regions show Cx40 localizes to gap junctions of Purkinje cells that display lower sarcomere content (F-Actin, red) (i.) and lack T-Tubules (WGA, green) (ii.) in contrast to surrounding cardiomyocytes, scale bar 20 μm. Cx40 specifically marks myocardial Purkinje fibers (PFs) diffusely distributed in the anterior left ventricular wall cross section, scale bar 2 mm (B).

**Online Video 1**

Purkinje fibers are distributed in a mesh-like network throughout the native porcine myocardium. Confocal z-stack of 28 images, z-step size 0.7 μm. Blue=DNA (Hoechst), Green=WGA, Red=F-Actin (phalloidin), White=CX40

**Online Video 2**

hESC-cardiomyocytes marked by slow skeletal troponin I (ssTnI) interact with connexin 40+ Purkinje fibers. Confocal z-stack of 56 images, z-step size 1 μm. Blue=DNA, Green= F-Actin, Red=ssTnI, White=CX40.

