## Supplementary figures and images for "Pharmacologic Therapy for Engraftment Arrhythmia Induced by Transplantation of Human Cardiomyocytes"

### Supplemental Figure 1

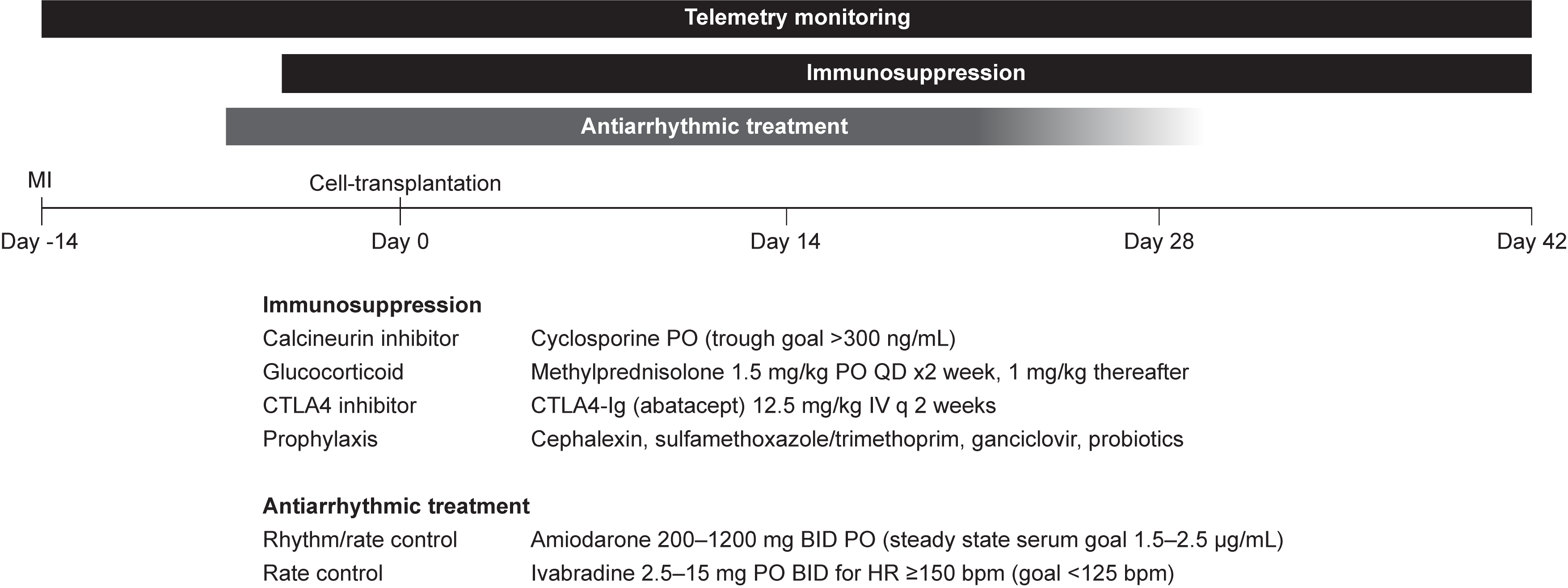

### Supplemental Figure 2

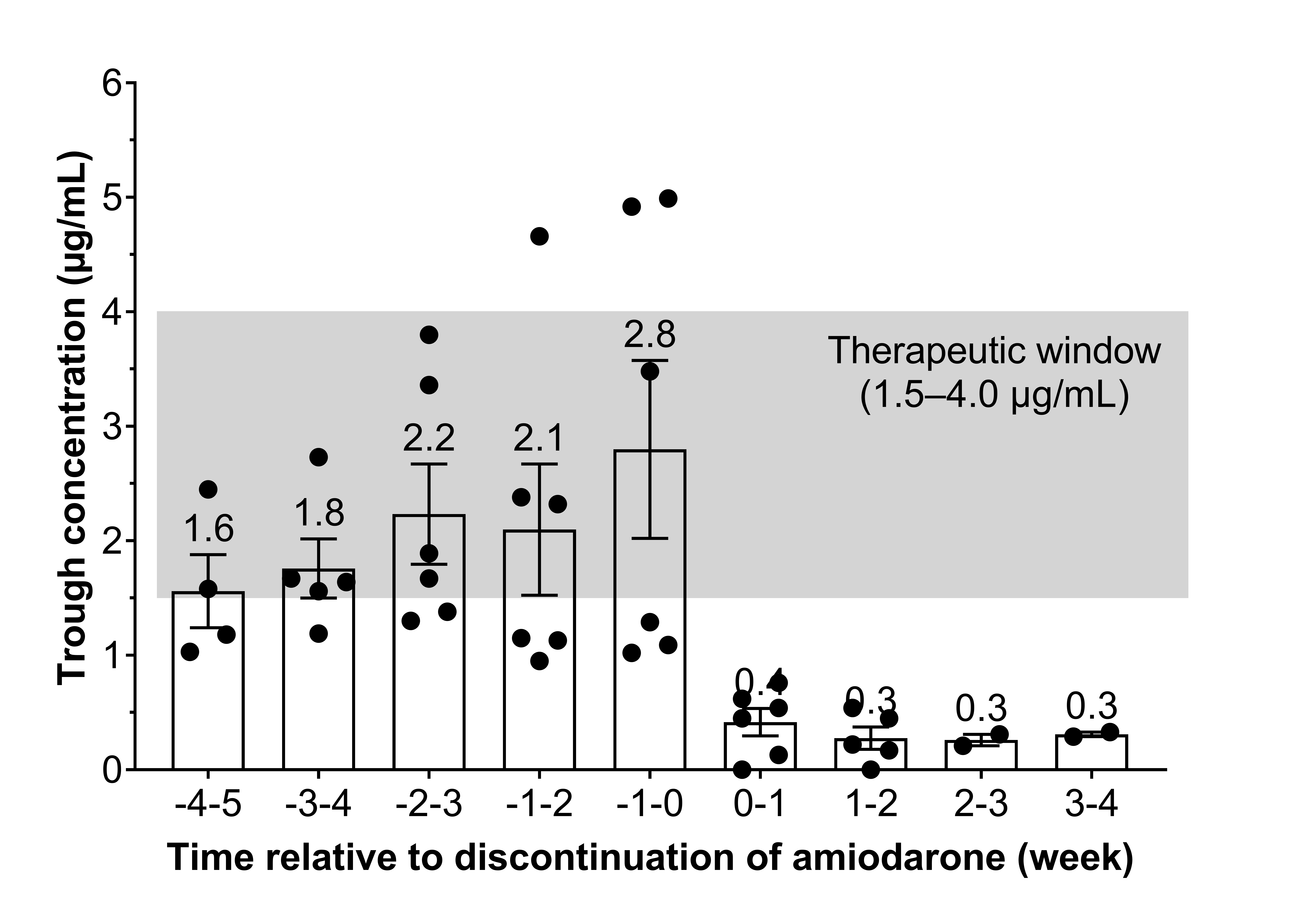

### Supplemental Figure 3

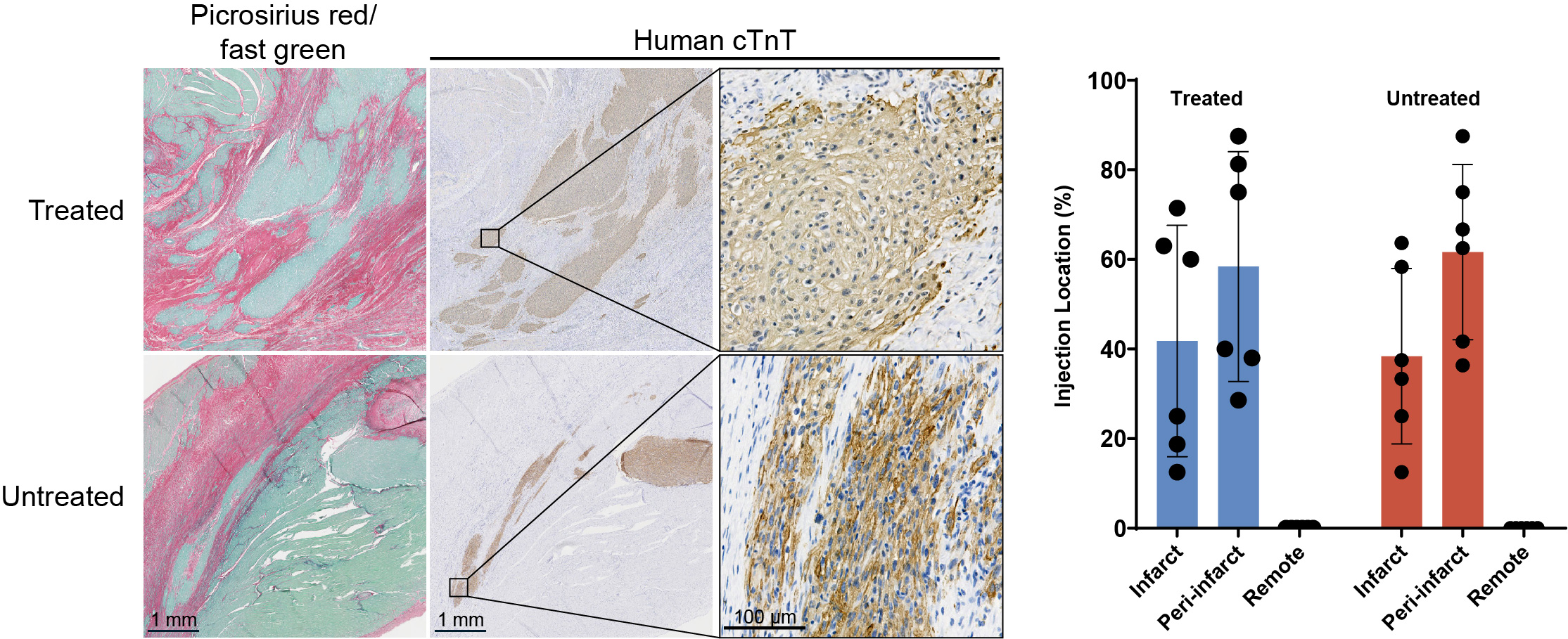

### Supplemental Figure 4

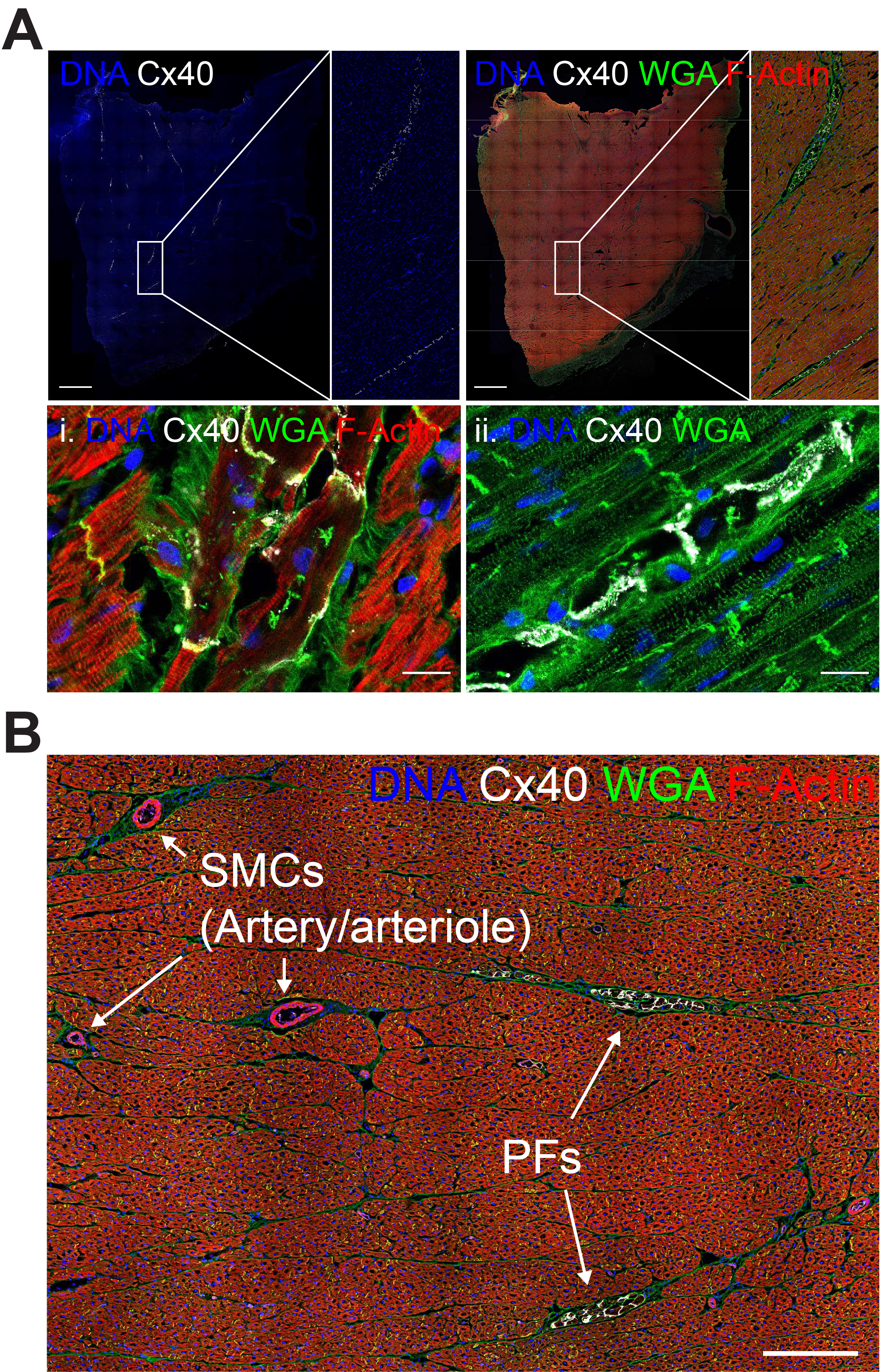
