## Supplemental Methods for "Pharmacologic Therapy for Engraftment Arrhythmia Induced by Transplantation of Human Cardiomyocytes"

### Online Methods

#### Animal subject care

All protocols were approved and conducted in accordance with the University of Washington (UW) Office of Animal Welfare and the Institutional Animal Care and Use Committee. Animals received ad libitum water and were fed twice a day (Lab Diet-5084 Laboratory Porcine Grower Diet). For surgical procedures, anesthesia was induced with a combination of intramuscular butorphanol, acepromazine and ketamine. Animals were intubated and mechanically ventilated using isoflurane and oxygen to maintain a surgical plane of anesthesia. Vital signs were monitored continuously throughout each procedure. All animals received subcutaneous Buprenorphine SR-Lab (ZooPharm) for post-operative analgesia and were euthanized by intravenous Euthasol (Virbac). All post-mortem examinations were performed by a blinded board-certified veterinary pathologist.

##### Porcine myocardial infarction model

Percutaneous ischemia/reperfusion injury was induced as previously described in NHP (12) with modification for the porcine model. A 5–8 cm incision was made in the femoral triangle and the femoral artery was exposed by blunt dissection. Prior to obtaining vascular access, heparin was administered to achieve therapeutic anticoagulation (activated coagulation time > 250 sec). A 5-French guidewire/introducer sheath system (Terumo Medical) was placed into the femoral artery and secured. Continuous ECG, invasive arterial blood pressure, pulse-oximetry and capnography were monitored throughout the procedure. Intravenous amiodarone 150 mg and lidocaine 100 mg were administered as single boluses prior to ischemia to minimize the risk of arrhythmia. Under fluoroscopic guidance (OEC 9800 Plus, GE Medical Systems), a 5-French Judkins right 2 or hockey stick guide catheter (Boston Scientific) was advanced into the ascending aorta to selectively engage the ostium of the left main coronary artery. Coronary angiography was performed using hand injections of contrast (Visipaque) and a 0.035” coronary guidewire (Runthrough NS Extra Floppy, Terumo Medical) was placed into the distal left anterior descending coronary artery (LAD). An appropriately sized angioplasty balloon catheter was then positioned into the mid-LAD distal to the first diagonal branch artery and inflated to the minimum pressure required for total obstruction of distal perfusion as confirmed by angiography. Ischemia was confirmed by ST-segment elevation on the ECG. Animals were maintained under anesthesia with ventilatory and hemodynamic support for 90 minutes, after which the balloon was deflated to restore distal perfusion, again confirmed by fluoroscopy and ECG. The animal was observed for reperfusion arrhythmias and externally cardioverted if ventricular fibrillation occurred. Prior to recovery, all animals received implantable telemetry units and central venous catheter placement. Briefly, the external jugular vein in the jugular furrow was exposed and a 5-French central venous catheter (Access Technologies) was inserted and tunneled out to the dorsal prescapular area. The telemetry transmitter (EMKA easyTEL+) was implanted in a subcutaneous pocket using the same incision in the jugular furrow, and subcutaneous leads were tunneled to capture the cardiac apex to base. The overall procedural mortality including the infarct was < 10%.

##### Immunosuppression therapy

All three cohorts received a three-drug immunosuppression regimen to prevent xenograft rejection as previously described with modification (12). For our initial regimen (subjects 1–6), five days prior to cell transplantation, oral cyclosporine A was started to maintain serum trough level of > 400 ng/mL (approximately 250–1000 mg twice daily) for duration of the study. Two days prior to transplantation, oral methylprednisolone was started at 3 mg/kg for two weeks then titrated down to 1.5 mg/kg for the remainder of the study. On the day of transplantation, Abatacept (CTLA4-Ig, Bristol-Myers Squibb) 12.5 mg/kg was administered intravenously and dosed every two weeks thereafter. Due to complications related to immunosuppression (principally porcine cytomegalovirus and Pneumocystis pneumonia), the cyclosporine A trough level was decreased to > 300 ng/mL and the methylprednisolone reduced to 1.0 mg/kg for subjects 7–19 without histologic evidence of rejection. Prophylactic oral cephalexin was administered for all subjects to prevent infection of the indwelling central venous catheter. Prophylactic sulfamethoxazole/trimethoprim was added after subject 3 developed Pneumocystis pneumonia. Prophylactic valganciclovir and probiotics were added after activation of endogenous porcine cytomegalovirus was found in subject 6.

#### Pilot antiarrhythmic screening in pig

Five infarcted pigs underwent hESC-CM transplantation with all exhibiting stereotypic EA. Subjects were administered multiple trials antiarrhythmics and observed for acute response by continuous ECG monitoring. Intravenous agents were delivered as a bolus dose over 2 minutes. Oral agents were administrated by direct observation in a minimum of apples, apple sauce or pumpkin puree with daily feeding and dosed daily for dose escalation. A washout period of at least three days was provided between agents. Amiodarone was administered as the last agent for testing given concern for prolonged half-life and elimination kinetics. All agents were tested in at least two subjects.

#### Amiodarone drug monitoring

Liquid chromatography–mass spectrometry was used to monitor steady state serum levels of amiodarone in the minipig model and guide oral dosing to ensure efficacy and avoid dose-related toxicity. A target serum level of 1.5–4.0 µg/mL was extrapolated from prior human pharmacokinetic studies (19,20). Elimination kinetics after discontinuation of oral amiodarone therapy were also studied by obtaining weekly trough concentrations in 6 subjects (6, 7, 8, 13, 14, 16) (Online Fig. 2).

#### Electrocardiography (ECG) analysis

Telemetric ECG was continuously monitored in real-time from the time of myocardial infarction to detect the primary endpoint of cardiac death or unstable EA. Automated quantification of heart rate and arrhythmia burden was performed offline by a board-certified cardiologist using the ecgAUTO 3.3.5.10 software package (EMKA Technologies). Arrhythmia was defined as an ectopic beat (e.g., premature ventricular contraction) or rhythm (e.g., idioventricular rhythm, ventricular tachycardia). EA was typically observed as sustained and non-sustained ventricular tachyarrhythmia of varying rates and morphologies but also included slow and narrow complex ectopic rhythms (Fig. 2). Heart rate and arrhythmia burden were quantified for two continuous minutes in each five-minute interval (40% of total rhythm was counted) and presented as daily averages.

#### Histologic analysis

Histological studies were carried out as detailed previously with modification (11,12). Briefly, paraformaldehyde-fixed hearts were dissected to remove the atria and right ventricle before short-axis cross-sections were cut at 2.5 mm intervals. The weights of the whole heart, left ventricle and each slice were obtained before further partition into tissue cassettes. The tissue then was processed, embedded in paraffin, and 4 µm sections were cut for staining. For morphometry, infarct regions were identified by picrosirius red staining; human graft was identified by anti-human cardiac troponin T (cTnT, Invitrogen, MA5-12960), stained using avidin-biotin reaction (ABC Kit, VectorLabs) followed by chromogenic detection via diaminobenzidine (Sigmafast, Sigma Life Science) (Online Fig. 3). The slides were digitized using a whole slide scanner (Nanozoomer, Hamamatsu), and the images were viewed and exported with NDP.view 2.6.13 (Hamamatsu). Areas of infarct and graft were analyzed using a custom-written algorithm in the ImageJ open source software platform (21). Briefly, after extracting images in TIFF format (22), the image foreground was segmented by a threshold derived from the distribution in brightness of its pixels, resulting in a binary mask that delineates the imaged tissue section. Subsequent color de-convolution by thresholding hue, brightness and saturation allowed segmentation of regions stained by Picro-Sirius Red stain or areas immunolabelled for human cardiac troponin-T. To separate scar from diffuse fibrosis, a cut-off for particle size was applied. Infarct size and graft size were calculated (percent area x block weight), summed for the entire ventricle, and expressed as a percentage of left ventricular mass or infarct mass, respectively. Please see the Online Methods for Purkinje fiber staining.

#### Purkinje fiber histology

For thin sections, tissue was cut and trimmed to 1 cm 🞪 1 cm 🞪 3 mm, snap frozen in isopentane, and embedded in OCT (TissueTek). 10 μm sections were immersed in 100% methanol at -20°C for 15 minutes and stained with standard immunofluorescence technique using stains described below. Images were acquired on a Leica SP8 confocal microscope.

For thick sectons, 1 cm 🞪 1 cm 🞪 3 mm pieces of tissue containing graft were incubated in 100% methanol at 20°C for 1 hour, rehydrated (80% methanol, 60% methanol, 0% methanol, diluted in PBS, 15-minute incubation at -20°C for each reagent). 150 μm sections were cut on a Leica VT1200s vibratome and stained with standard immunofluorescence technique using stains described below. Stained sections were then cleared using BABB as previously reported (37), and imaged on a Leica SP8 confocal microscope with 1 μm z-step increments.

#### Purkinje fiber staining

Sections were stained the following reagents: Hoechst 33342 (DNA, Thermo Fisher Scientific, #62249), Wheat germ agglutinin-Oregon Green (WGA, Thermo Fisher Scientific, #W6748), Phalloidin-647 (F-Actin, Thermo Fisher Scientific, #A22287), anti-Connexin 40 (Cx40, Alpha Diagnostics, #CXN40A), or anti-slow skeletal troponin I (ss-TnI, Novus, #NBP2-46170) with one of two anti-rabbit secondary antibodies (Alexa Fluor 555/647, Thermo Fisher Scientific, #A-31570/A-31573).
