## Supplemental Table for "Pharmacologic Therapy for Engraftment Arrhythmia Induced by Transplantation of Human Cardiomyocytes"

### Supplemental Materials

#### Supplemental Table 1

| **Lidocaine (Ib)** | 100 mg IV | Modest HR effect, rare cardioversion |
| --- | --- | --- |
| **Flecainide (Ic)** | 2 mg/kg PO, 4 mg/kg PO, 6 mg/kg PO, 10 mg/kg PO | No response on HR or EA burden |
| **Propafenone (Ic)** | 1 mg/kg IV, 2 mg/kg IV, 3 mg/kg IV | No response on HR, transient cardioversion* |
| **Amiodarone (III)** | 150 mg IV | Modest HR effect, frequent cardioversion |
| **Sotalol (III)** | 1 mg/kg PO, 2 mg/kg PO, 4 mg/kg PO | No response on HR or EA burden |
| **Metoprolol (β_1_AR)** | 5 mg IV, 25 mg PO BID, 50 mg PO BID, 75 mg PO BID | Moderate HR effect (IV only), no response on EA burden |
| **Ivabradine (I_f_)** | 2.5 mg PO, 5 mg PO BID, 10 mg PO BID, 15 mg BID; 1 mg/kg IV, 2 mg/kg IV | Robust dose-dependent HR effect (PO only)**, no response on EA burden |
| * Severe nausea/emesis observed at therapeutic doses, limiting clinical utility | | |
| ** Severe bradycardia | | |
| Abbreviations: β_1_AR, β1-adrenergic receptor; BID, twice daily; HR, heart rate; EA, engraftment arrythmia; I_f_, funny current; PO, oral; VT, ventricular tachycardia | | |
